# Integrity of the Uncinate Fasciculus Predicts Emotional Pattern Separation-Related fMRI Signals in the Hippocampal Dentate and CA3

**DOI:** 10.1101/858258

**Authors:** Steven J. Granger, Stephanie L. Leal, John T. Janecek, Liv McMillan, Hal Stern, Michael A. Yassa

## Abstract

Alterations in white matter integrity have been demonstrated in a number of psychiatric disorders that involve disruptions in emotional processing. One such pathway – the uncinate fasciculus (UF) – connects the orbitofrontal cortex (OFC) to the medial temporal lobes (MTL) and has been associated with early life adversity, maltreatment, anxiety, and depression. While it is purported to play a role in episodic memory and discrimination, its exact function remains poorly understood. We have previously described the role of the amygdala and dentate (DG)/CA3 fields of the hippocampus in the mnemonic discrimination of emotional experiences (i.e. emotional pattern separation). However, how this computation may be modulated by connectivity between the medial temporal lobes and the orbitofrontal cortex remains unknown. Here we ask the question of whether the uncinate fasciculus plays a role in influencing MTL subregional activity during emotional pattern separation. By combining diffusion imaging with high-resolution functional MRI, we found that reduced integrity of the UF is related to higher activation in the DG/CA3 subregions of the hippocampus during an emotional pattern separation task. We additionally report that higher levels of DG/CA3 activity are associated with poorer memory performance, suggesting that hyperexcitability in this network (which may be driven by CA3 recurrent collaterals) is associated with memory errors and that the UF may allow the OFC to exert inhibitory control on this network and improve discrimination of emotional experiences. This work provides novel mechanistic insight into the role of prefrontal interactions with the MTL, particularly in the context of emotional memory.

## Introduction

Episodic memory – memory for facts and events that shape our lives – involves contributions from the medial temporal lobe (MTL) as well as the prefrontal cortex (PFC) as well as their interactions (Eichenbaum, 2017; Preston & Eichenbaum, 2013). While a considerable imaging literature in humans has focused on the hippocampus, relatively few studies have examined the role of white matter pathways connecting the MTL to the PFC in supporting episodic memory computations. Studies in rodents have been hindered by the major species differences in PFC anatomy as well as the absence of certain anatomical connections that are present in primates.

One such anatomical connection, the uncinate fasciculus (UF), is a major white matter bundle connecting the MTL to the orbitofrontal cortex (OFC) in primates. This particular connection forms an “anterior pathway” by which the MTL can communicate with the PFC and has been thought of as a top-down regulator involved in fear-related memory processes (Marek et al., 2018). In humans, deficits in the UF have been implicated in psychiatric disorders characterized by emotional dysregulation such as major depression and anxiety, as well as in early life stress and maltreatment (Taylor et al., 2007; Zhang et al., 2012; Eluvathingal et al., 2006; Ho et al., 2017; Hanson et al., 2015). Although often described as innervating the hippocampus, human dissection and axonal tracing studies in primates have indicated that this bundle more accurately innervates the basolateral amygdala (BLA) and the entorhinal cortices of the MTL (Ebeling and Von Cramon, 1992; Von der Hide et al., 2013; Thiebaut de Schotten et al., 2012). Further, the UF has distinct projections to a number of prefrontal areas in particular the OFC (**Fig. 1**).

**Fig 1.**
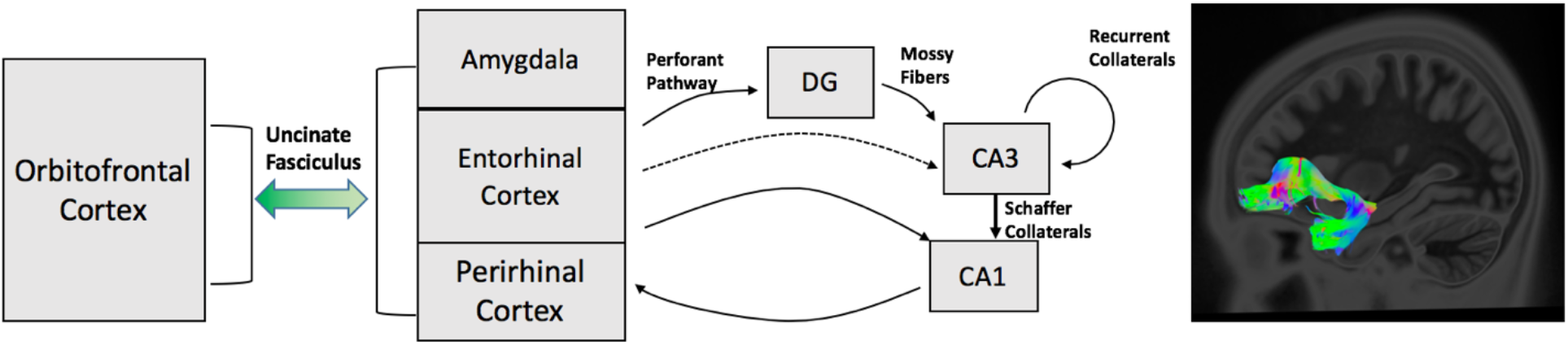
Circuitry of orbitofrontal connectivity with the medial temporal lobe subfields via the uncinate fasciculus. The uncinate innervates both the lateral amygdala and cortices of the MTL. The cortices of the MTL feed into the hippocampal subfields of DG, CA3, and CA1 via multiple pathways one of which includes the perforant path. The perforant path feeds into the DG/CA3 subfields where the neuronal computation known as pattern separation is thought to occur. In this manner, the uncinate fasciculus is a possible direct route of inhibitory information to these regions.

While the exact functional role of the UF remains unclear, it is thought to be involved in episodic memory, language, and socio-emotional processing (Von Der Heide et al., 2013). The UF is also hypothesized to play a role in adjudicating among competing episodic memory representations at retrieval, a hypothesis which has gained some recent empirical support (Alm et al., 2016). Competition among memory traces during retrieval could be mediated, at least in part, by pattern separation. This computation, as first suggested by Marr (Marr, 1971), is defined as the neural computation used to orthogonalize overlapping similar stimuli, a process that particularly involves the dentate gyrus and CA3 regions of the hippocampus (Yassa & Stark, 2011; Leal and Yassa 2018). Given this prior work, we reasoned that tasks assessing pattern separation would be particularly well-suited to understanding the impact of the UF connection on episodic memory.

High-resolution fMRI studies have noted pattern-separation related activity in the DG/CA3 region during incidental encoding of similar lure items (Bakker et al. 2008; Lacy et al. 2011) as well as explicit performance of mnemonic discrimination tasks (Yassa et al. 2010a, 2010b; Reagh et al. 2018). These tasks require participants to determine during retrieval if items highly similar to the previously encoded items are “new” or “old”. This challenging discrimination procedure requires suppression of interference from competing memory representations. In the context of emotional memories, this network extends to also include the amygdala (Leal et al. 2014b). We recently showed using intracranial recordings from presurgical epilepsy patients that alpha-band mediated unidirectional influence of the amygdala on the DG/CA3 during an emotional mnemonic discrimination task is associated with overgeneralization errors (Zheng et al. 2018). This suggests that MTL computations can be biased to discriminate or generalize across similar experience but the role of the PFC or MTL-PFC connections in setting this bias is unclear.

As one of the major structural pathways connecting the MTL and the PFC, the UF is a likely contributor to the process of emotional memory discrimination. However, its role in this process has never before been investigated. Using multimodal neuroimaging, we tested the hypothesis that the UF regulates mnemonic discrimination specifically for emotional items by altering BOLD fMRI activity in the DG/CA3 activity, which in turn would predict emotional discrimination performance. To assess the structural role of the UF, we implement a novel way to anatomically track the pathway using non-tensor-based or “model free” diffusion weighted imaging and deterministic tractography (Yeh et al., 2013). We quantified the integrity of the UF by utilizing a model-free diffusion metric (normalized quantitative anisotropy – nQA) which is designed to capture the degree of diffusion among primary fiber orientations accounting for partial volume effects indicative of the tensor model (Yeh et al., 2013). To assess the functional role of the DG/CA3 region of the hippocampus, we used high-resolution (1.5mm isotropic) BOLD fMRI capable of resolving hippocampal subfields, while participants performed the emotional discrimination (i.e. pattern separation) task (**Fig. 2**). The outcome measure was the “lure discrimination index”, a measure of how well participants are able to discriminate among similar emotional and neutral scenes, corrected for response bias.

**Fig 2.**
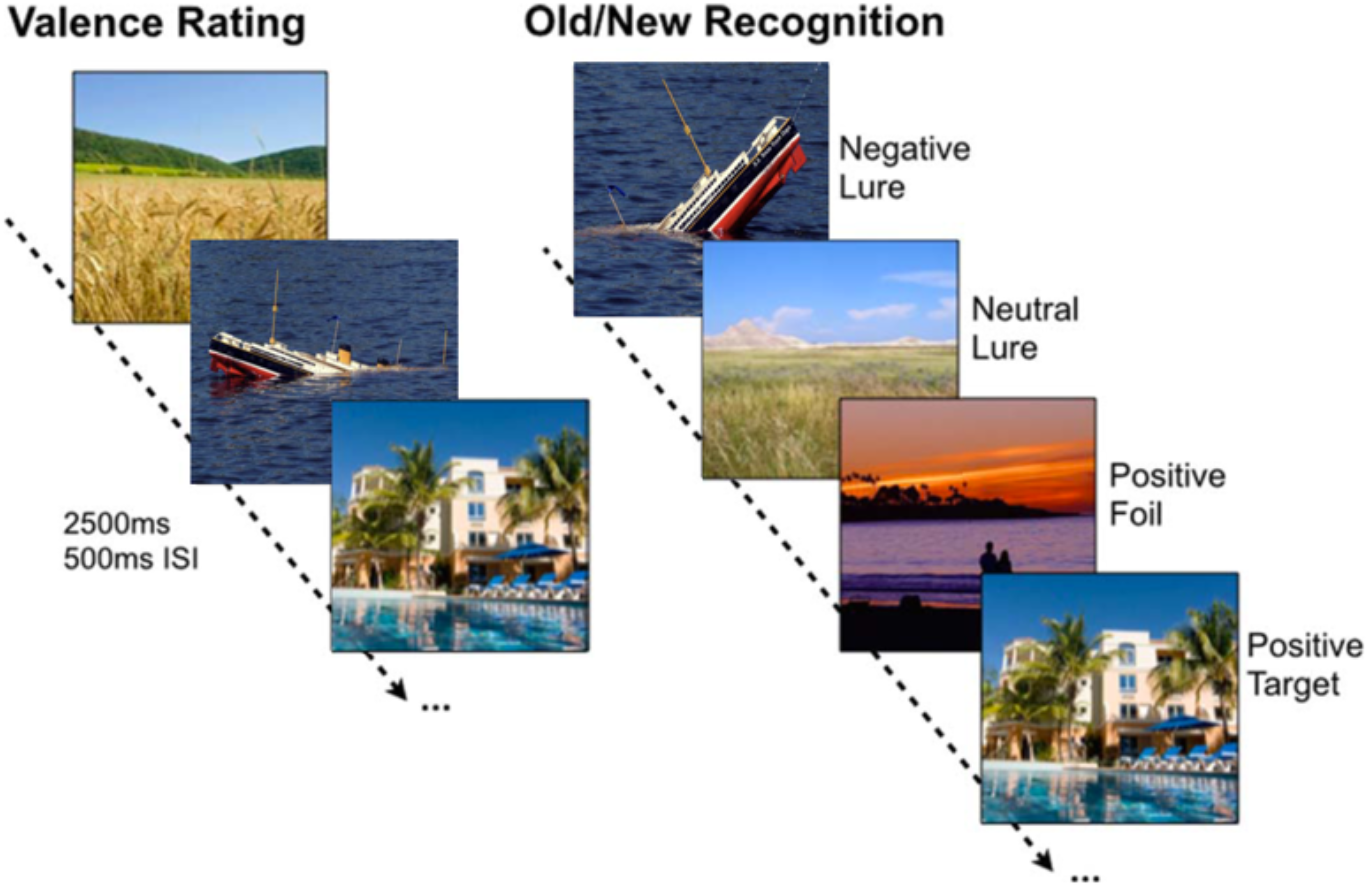
Schematic of the emotional pattern separation task. BOLD activity from DG/CA3, CA1, and amygdala. The left side of the image shows the incidental encoding phase of the task where participants are asked to rate images as “negative”, “neutral”, or “positive”. On the right side of the image is the memory portion of the task. This phase, known as the Old/New Recognition phase shows participants several variants of previously seen images (lures), as well as repeated images (repeats), and never before seen images (foils). Subjects are asked to make “old” or “new” judgements during the trial and response times are recorded.

## Results

### Uncinate Integrity Predicts DG/CA3 Activity During Emotional Lure

To test the hypothesis that UF structural integrity is associated with MTL subregional activation profiles during discrimination of highly similar lures, we used normalized quantitative anisotropy (nQA) to quantify the UF and BOLD fMRI activity in the DG/CA3 region during correct discrimination as our variables of interest. Here we separately assessed the magnitude of this relationship (between UF integrity and DG/CA3 activity) in highly similar negative, neutral, and positive correct discrimination trials. We focused on high-similarity lures as prior work suggested that DG/CA3 activity is particularly sensitive to this condition (Leal et al. 2014b). We found a significant negative correlation between left UF nQA and left DG/CA3 activity for highly similar negative (*n* = 27, Pearson *r* = −0.4067, *P* = 0.0353) and positive (*n* = 27, Pearson *r* = −0.4385, *P* = 0.0221) but not neutral (*n* = 27, Pearson *r* = 0.1206, *P* = 0.5491) CRs (**Fig 3a-c**).

**Fig 3.**
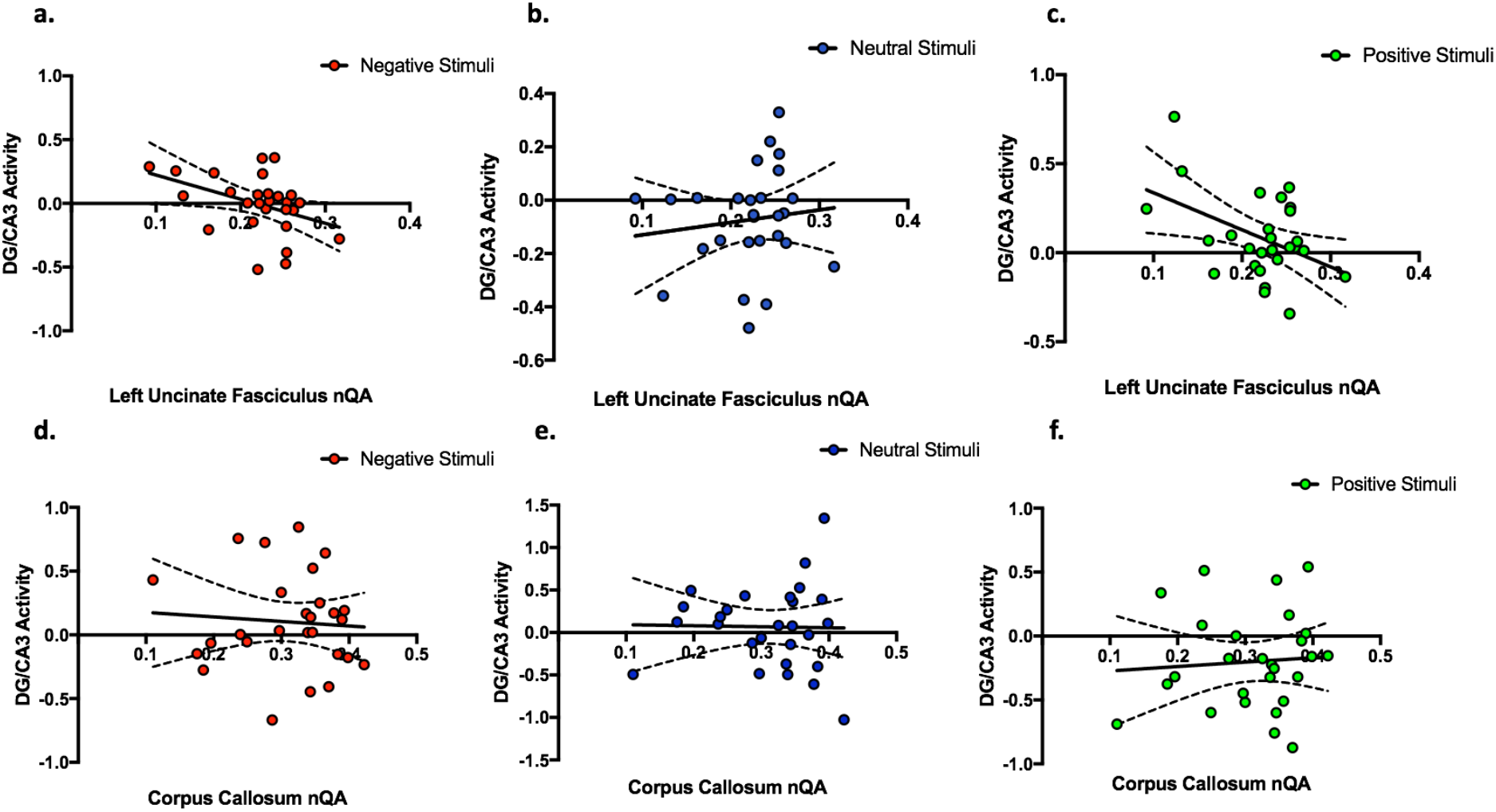
Comparison between structural measures predicting functional measures Left uncinate fasciculus nQA predicting left DG/CA3 BOLD activity during highly similar lure correct rejections. (a-c) Shows the left uncinate fasciculus nQA predicting DG/CA3 BOLD response during the correct discrimination of (b) negative (*r* = −0.4067, *P* = 0.0353) and (d) positive (*r* = −0.4385, *P* = 0.0221) highly similar lure items but not (c) neutral (*r* = 0.1206, *P* = 0.5491). (e) Shows the corpus callosum ROI as our negative control. (f-h) No relationship between corpus callosum nQA and average left and right DG/CA3 activity during the correct discrimination of (f) negative, (g) neutral, and (h) positive highly similar lure items.

We then tested if this the relationship between UF integrity and DG/CA3 activity during lure discrimination was specific to the DG/CA3 subfield of the hippocampus. We found that that left UF nQA did not predict with the responses of the left CA1 subfield. Additionally, these effects seem specific to the left hemisphere. Right UF integrity did not predict right DG/CA3 nor CA1 activity during highly similar lure discrimination for any valence type. All p-values were above .05.

To further test the specificity of UF nQA predicting DG/CA3 activity during highly similar emotional lure discrimination, we tested if UF nQA also predicted DG/CA3 activity during low similarity lure correct rejection. Left UF nQA did not predict DG/CA3 activity for negative (n = 27, Pearson *r* = 0.0838, *P* = 0.6775), neutral (n = 27, Pearson *r* = 0.214, *P* = 0.283), or positive (n = 27, Pearson *r* = 0.152, *P* = 0.4487) low similarity correct rejections.

As the UF has been previously implicated in psychiatric disorders characterized by emotional disturbances, we asked if this effect could be related to symptoms of depression. Using the Beck Depression Inventory II (BDI-II) we quantified depressive symptoms in all participants and included the overall score in a regression model (BDI-II and UF nQA predicting DG/CA3 activity). We found that the relationship between left UF nQA and left DG/CA3 activity for highly similar negative lure correct discrimination trials remained significant (*B* = −1.8026, p-value= 0.0481) after accounting for BDI-II scores (B = −0.002863, p-value = 0.4555). Similarly, the relationship between left UF nQA and left DG/CA3 activity for highly similarly positive items remained significant (*B* = - 1.941544, p-value 0.03220) after accounting for BDI-II scores (*B* = −0.004507, p-value = 0.23795).

To determine if this relationship was specific to the UF or is a general feature of white matter connectivity, we tested the relationship between general connectivity and DG/CA3 activity (using the corpus callosum as a control pathway). We tested the hypothesis that the corpus callosum nQA predicted averaged DG/CA3 response during high similarity negative, positive and neutral lures. We found that none of the relationships were statistically significant (**Fig 3d-f**). Together, these data suggest that *lower* integrity of the UF is associated with higher levels of activity in the DG/CA3 during correct discrimination, suggesting a possible role in inhibiting or limiting the excitability of this subregion, and that this effect is specific to the UF and is not a general characteristic of other white matter.

### DG/CA3 Functional Activity Predicts Lure Discrimination

Next, we assessed the relationship between DG/CA3 subfield activity during discrimination of highly similar lure was associated with overall performance on those trial types. Past work in our group and others has suggested that higher levels of activity in the DG/CA3 region is associated with overgeneralization errors. We found a significant negative correlation between left DG/CA3 activity during correct discrimination of negative stimuli and the lure discrimination index on the same trials (n = 27, Pearson *r* = −0.4205, p-value = 0.029, **Fig. 4**). We once again assessed whether depressive symptoms impacted this relationship by including BDI-II score as a regressor and found that that the relationship between left DG/CA3 activity and negative LDI remained significant (*B* = −0.313636455, p-value = 0.0399) accounting for depressive symptoms (*B* = 0.0008924, p-value = 0.7633).

**Fig 4.**
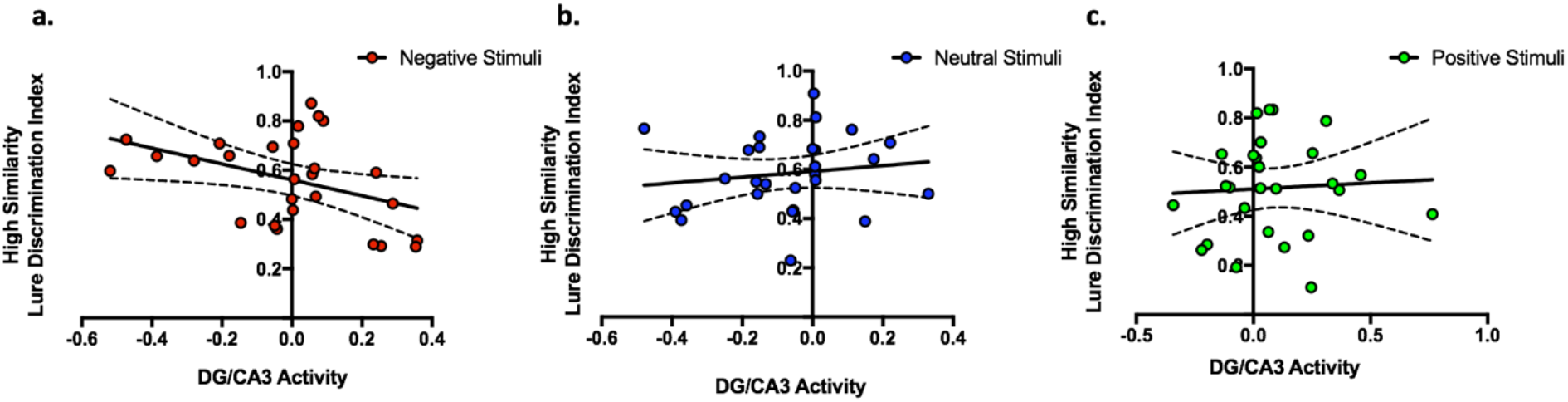
Relationship between DG/CA3 BOLD response and lure discrimination behavior. Left DG/CA3 activity during high similarity CRs predicting lure discrimination for (a) negative (*r* = −0.4205, p-value = 0.029), but not (b) neutral or (c) positive stimuli.

### DG/CA3 Functional Activity Mediates the Association Between the Uncinate Fasciculus and Lure Discrimination

Given the relationship between the UF and DG/CA3 activity, as well as the relationship between DG/CA3 activity and lure discrimination performance, we asked whether UF integrity directly predicts lure discrimination. We found no significant correlations between left UF nQA and LDI of highly similar negative (n = 27, Pearson *r* = 0.2359, p-value = 0.2362), neutral (n = 27, *r* = −0.1288, p-value = 0.5220), or positive items (n = 27, Pearson *r* = 0.2423, p-value = 0.2233). We also found no significant correlations between right UF nQA and LDI of highly similar negative (n = 27, Pearson *r* = 0.2918, p-value = 0.1397), neutral (n = 27, Pearson *r* = −0.02436, p-value = 0.9040), or positive items (n = 27, Pearson *r* = 0.2247, p-value = 0.2598). We then asked if perhaps the relationship between UF integrity and behavioral performance may be mediated by the functional MRI signal noted in the DG/CA3, since this signal correlates significantly with both UF integrity and with discrimination performance.

We implemented a mediation analysis testing whether DG/CA3 activity mediates the relationship between UF nQA and lure discrimination for highly similar negative lure items. We found a marginally significant average causal mediation effect (ACME) of DG/CA3 activity during CRs of highly similar negative items predicting negative LDI (keeping uncinate nQA constant) (β = 0.5638, p-value = 0.079). Additionally, neither the average direct effect (β = 0.2772, p-value = 0.60), total effect (β = 0.108, p-value = 0.8409), or proportions mediated (β = 0.6704, p-value = 0.163) were significant (**Fig. 5**). This analysis suggests the possibility that UF may serve as a pathway to regulate DG/CA3 activity and that DG/CA3 is ultimately what influences discrimination behavior, although given the marginally significant findings, this result should be interpreted with caution.

**Fig 5.**
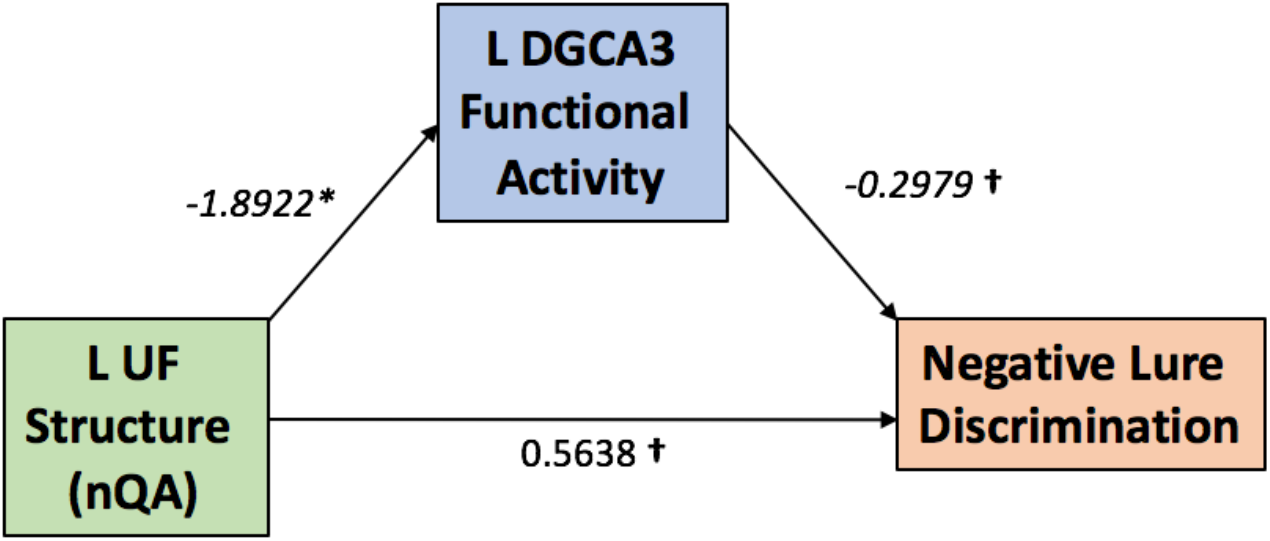
Schematic summary of mediation results. Here we show decreases in the static measure of uncinate fasciculus integrity are associated with increased activity in the DG/CA3 subfield of the hippocampus and better discrimination performance on negative lures. We provide marginal evidence that UF structure exerts its influence on behavior through altering DG/CA3 activity (β = 0.5638, *P* = 0.079) possibly as a pathway involved in top-down regulation of hippocampal processes in episodic memory function. *Significant at alpha=0.05. †Marginally significant.

## Discussion

In this study, we tested the hypothesis that the integrity of the uncinate fasciculus is associated with medial temporal lobe dynamics during an emotional pattern separation task. This stems from the observation that prefrontal cortex modulation of MTL signaling is implicated in memory processing (Jones & Wilson, 2005; Kim et al., 2011; Brincat & Miller, 2015; Kesteren et al., 2010), however, the exact pathways by which this modulation occurs in humans have remained elusive.

Accounting for depressive symptoms, we found that in the left hemisphere, lower uncinate fasciculus integrity predicted greater activation of the DG/CA3 subfield of the hippocampus during emotional (positive and negative) highly similar correct discrimination trials. To our knowledge, this is the first demonstration that differences in uncinate fasciculus integrity may be associated with alterations of functional activity in the DG/CA3 subfields of the hippocampus during discrimination of similar information. These findings provide evidence for a potential link between uncinate integrity and hippocampal activity during emotional conditions.

In addition to providing a link between uncinate structure and DG/CA3 activity, we found that greater left DG/CA3 activity during highly similar negative correct rejections was related to poorer memory performance for highly similar negative items (i.e. higher rate of false alarms). Although at first this result seems surprising, given what is known about the role of DG/CA3 in pattern separation, a finding of increased activity in DG/CA3 linked to poor performance has been previously reported in older age (Yassa et al., 2011; Sinha et al., 2018). An interesting possibility is that the increased excitability is related to CA3’s recurrent collateral network which is thought to play a role in pattern completion (or overgeneralization in the case of a discrimination task). While the noted activity occurred during the correct trials, pattern completion is assumed to occur also during correct lure discrimination (i.e. “recall to reject”). Here, much like the data in older adults, higher levels of DG/CA3 activity during correct lure discrimination is linked to poorer overall memory performance at the individual subject level. Overall, this suggests that hyperactivity in this subfield is associated with worse cognitive outcomes. That said, other work has shown that a within-subject increase in DG/CA3 activity as a function of mild aerobic exercise is associated with enhanced discrimination (Suwabe et al. 2018), however this finding also was associated with increased functional connectivity with regions involved in high precision recollection of memories (e.g. restrosplenial cortex, angular gyrus). Given this evidence, it is possible that there is an inverted U-shaped dose-response relationship where activity in DG/CA3 needs to be held in balance, with too much or too little activity being associated with worse outcomes. This particular topic should be subject to further investigation.

Our mediation model suggests that while uncinate integrity was not a robust direct predictor of lure discrimination behavior, its impact on behavior may be mediated by DG/CA3 activity. The lack of correlation between uncinate integrity and discrimination performance is consistent with other results from Bennet et al., (2015), who found no link between uncinate integrity and object discrimination performance. Instead, the authors found a relationship between the fornix and performance, a relationship that was consistent across young and older adults. The possibility that the uncinate’s impact on behavior is mediated by functional signals in the DG/CA3 is consistent with a top-down modulatory control from the PFC to the MTL during discrimination.

Although we tested a focused hypothesis about the uncinate fasciculus and ensured the specificity of the findings by comparison against a negative control pathway (the corpus callosum), it is possible that other pathways might influence these emotional memory processes. In general, the anatomical connectivity of the uncinate fasciculus, cingulum, and fornix bundles and the relative contributions to MTL-PFC connectivity in humans remains poorly understood. The three-dimensional structures of these pathways have not been extensively resolved in humans or nonhuman primates *in vivo*. It is possible that information flow is segregated according to information content and valence. For example, signaling involving emotional items could be reliant on OFC integration and thus be communicated via the uncinate fasciculus, whereas signaling involving non-emotional items could be reliant on mPFC (or other PFC region) integration and thus be communicated via the cingulum or fornix. This may be too simplified, as it is not currently known whether these pathways innervate overlapping subdivisions of the PFC, and it is unclear what the functional consequences of these dissociations may be. While speculative, this division of labor should be subject to further investigation.

One issue for our work and others is the difficulty of even modern tractography algorithms and data acquisition schemes to accurately and non-invasively model even the largest white matter bundles with sufficient accuracy in humans (Maier-Hein et al., 2017). Tractography of the uncinate fasciculus is a particularly challenging endeavor due to its branching and turning patterns and the possibility for false continuation medially toward the striatum and posteriorly along the inferior fronto-occipital fasciculus and inferior longitudinal fasciculus.

Our use of more sophisticated tractography algorithms that account for multiple fiber orientations within each voxel might aid in the delineation of this morphologically complex fiber. For example, we used normalized quantitative anisotropy, a measure that resolves multiple fiber orientations within a voxel and appropriately accounts for the isotropic component of diffusion, which addresses partial volume confounds. We combined this approach with anatomical regions of avoidance consisting of three intersecting planes and a sphere over the striatal area to ultimately generate a highly reproducible scheme for tractography of the uncinate fasciculus while avoiding common false continuations with respect to anatomy.

In this study, we have shown that the decreased UF integrity is associated with greater activity of DG/CA3 during emotional lure discrimination and that greater DG/CA3 activity is associated with poor performance on an emotional pattern however, this remains speculative and subject to further investigation. separation task. Our interpretation of these results is that the UF mediates important top-down control from the OFC to the MTL which is critical for processing of emotional memories, but that the relationship between UF integrity and behavior is mediated by functional activation of the hippocampal network. This suggests that UF integrity may be a marker for emotional and cognitive health, which is consistent with prior work implicating the UF in emotional dysregulation conditions including major depressive disorder and anxiety (Zhang et al., 2012), as well as work linking UF integrity with early life stress and maltreatment (Eluvathingal et al., 2000).

Future studies could examine whether the reported functional and structural findings manifest beyond emotional discrimination tasks or if there are changes in the relationship in UF’s structure and function in the context of development and/or aging.

## Materials and Methods

### Subjects

Twenty-seven subjects (12 male) were recruited from Johns Hopkins University and received monetary compensation for their participation. Informed consent was given by all participants and all procedures approved by the Johns Hopkins University Institutional Review Board. Subjects were administered a neuropsychological battery which is summarized in **Table 1**. All participants were screened against major medical or psychiatric morbidities, substance abuse history, and additional criteria of MRI contraindications like metal in the body. Results of the fMRI analysis of this dataset were previously published in Leal et al. 2014b.

**Table 1.** Participant Demographics and Neuropsychological Results

| Variable | Mean (± SD) |
| --- | --- |
| Age | 21 ± 3.38 |
| Beck Depression Inventory-II | 10.55 ± 11.48 |
| RAVLT Immediate Recall | 12.59 ± 1.97 |
| RAVLT Delayed Recall | 11.63 ± 3.44 |
| Digit Span Forward | 11.96 ± 1.85 |
| Digit Span Backward | 8.29 ± 2.03 |
| Mini Mental State Exam | 29.22 ± 0.97 |
| Trails Making Test A | 19.58 ± 5.31 |
| Trails Making Test B | 45.59 ± 11.91 |

### Imaging Data Collection

All data were collected on a 3-Tesla Philips scanner. We collected an ultrahigh-resolution structural MPRAGE scan that we developed for accurate delineation of hippocampal subfields and high-resolution diffeomorphic alignment (0.55 mm isotropic resolution; 273 sagittal slices, field of view = 240 x 240 mm, flip angle = 9, TR/TE = 13/5.9 ms, matrix size = 448 x 448, inversion pulse TI = 1110 ms). SENSE parallel imaging was used in two directions (2 x 1.5). The SAR (<10%) and PNS (<75%) were within required limits based on the scanner-calculated values. Task-activated functional data were collected and processed in the same manner as Leal et al., 2014b. Briefly, these data were collected with a high-speed EPI single-shot pulse sequence (1.5mm isotropic, 19 oblique axial slices parallel to the principal axis of the hippocampus, field of view = 96 x 96 mm, flip angle= 70, SENSE parallel reduction factor = 2, TR/TE = 1500/30 ms, matrix size = 64 x 64. Diffusion weighted imaging were collected with an echo planar imaging sequence. The data were 3 mm isotropic, TR/TE = 6800/67 ms, 65 slices, 32 non-collinear directions, one b0, b-value = 700 s/mm2. Diffusion MR data were collected at the time of the task-data.

### Emotional Pattern Separation Task Description

The emotional pattern separation task consists of an incidental encoding phase where subjects are instructed to rate the valence of each image from positive, negative, or neutral based on a priori ratings outlined in Leal et al., 2014a. Subjects were later tested during the “Old/New Recognition” portion of the task. In this portion, foils (new images), targets (same images), and high and low similarity lures were presented. Subjects were asked to make “Old” or “New” recognition judgments (*Figure 2*). In this case, correctly identifying that a lure item was “New” is termed as a Correct Rejection (CR) and falsely claiming that a lure item is “Old” would be termed a False Alarm (FA). We assessed behavioral discrimination on this task by quantifying a lure discrimination index (LDI) which measures performance on the task accounting for response bias: P(“New”|Lure) – P(“New”|Target) or lure correct rejections minus target misses. This was calculated for high and low similarity lures across the three valence types. LDI of highly similar lure items was chosen in this case because of the UFs’ purported role in adjudicating between similar items during retrieval and it also accounts for response biases unlike other performance measures like correct rejection rate.

### Functional MRI Image Analysis

Task data were analyzed using the open source package Analysis of Functional Neuroimages (AFNI) in the manner described in Leal et al., 2014b (Cox, 1996). Briefly, images were corrected for slice timing and subject motion censoring motion of 3° of rotation or 2mm translocation in any direction relative to prior acquisition. Upon processing for motion we then registered our functional images to the structural (MPRAGE) using AFNI/s 3dAllineate program. We use Advanced Normalization Tools (Avants et al., 2011) which implements a robust diffeomorphic algorithm to warp structural scans to a common template based on the entire sample. The same transformations were then applied to the functional data. Finally, behavioral vectors were created based on trial type orthogonalizing trials unique in emotion, similarity, and behavioral decision. These vectors there then used in a deconvolution approach based on multiple linear regression and the resulting fit coefficients (betas) estimated activity versus novel foils as an implicit baseline for a given time points and trial type in a voxel was calculated. We used the sum of the fit coefficients over an expected hemodynamic response of 3-12 seconds after trial onset as the model’s estimate of response to each trial type as described in Leal et al., 2014b. Finally, fit coefficients were extracted using a region of interest approach for the DG/CA3 and CA1 subfields of the hippocampus according to Leal et al., 2014b.

### Diffusion Weighted Imaging Analysis

Diffusion data were processed for eddy correction using FSLs *eddy_correction* and analyzed using DSI-Studio (http://dsi-studio.labsolver.org). Individual subject data were then processed using DSI-Studios Q-Spin Diffeomorphic Reconstruction (QSDR) method which calculates the orientational distribution of the density of diffusing water in MNI space (Yeh & Tseng, 2011). All subjects met the criterion of fitting to the template space by obtaining an R-squared value of greater than 0.63 as suggested by the software developers.

QSDR was chosen as the reconstruction method in order to overcome several limitations of the tensor model. First, QSDR provides reconstruction in a template space allowing the creating of standardized “regions-of-avoidance” to filter out known false-projecting fibers. Second, QSDR allowed for the computation of orientation distribution functions rather than tensor-based computation of diffusion signals. This is important because the orientation distribution functions calculated with this method are presumed to resolve partial volume fractions, model crossing-fibers, and presumably result in more accurate deterministic tractography (Yeh et al., 2013). Our tractography protocol included a custom region-of-avoidance by merging three intersecting planes (one axial, one coronal, one sagittal) with a sphere over the striatal and nucleus accumbens complex (**Fig. 6**).

**Fig 6.**
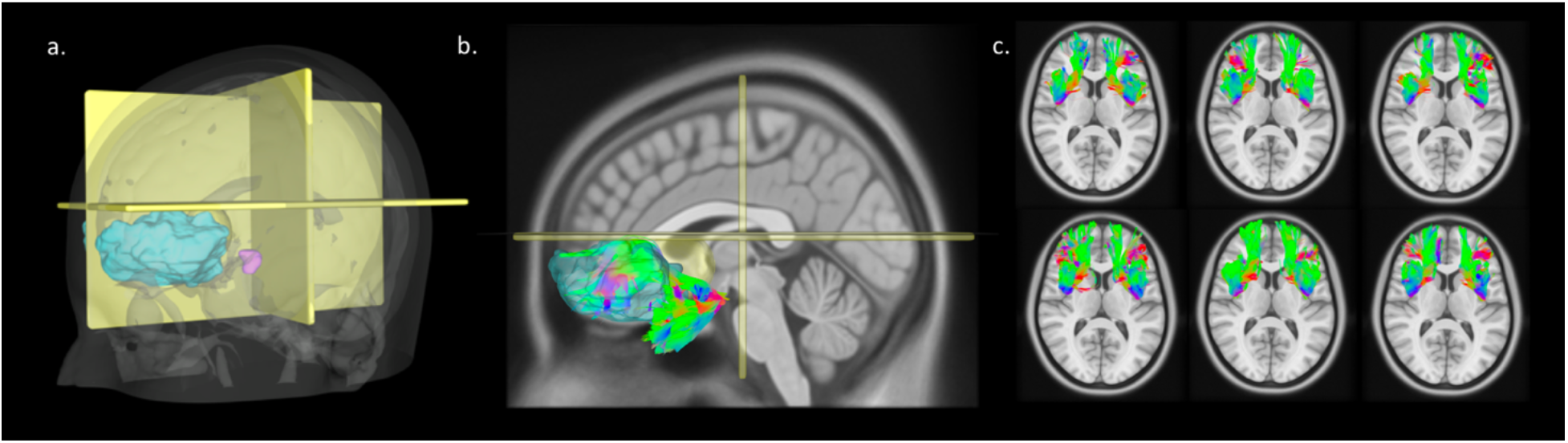
In-vivo dissection of the uncinate fasciculus derived from model-free QSDR and deterministic tractography in the template space. (a) The “region-of-avoidance” is depicted in yellow, uncinate fasciculus “seed region” depicted in purple, and “end” region in blue containing merged Brodmanns areas 11 and 47. (b) The resulting UF tractography for the left hemisphere. The tractography shows branching in the prefrontal cortex region as anatomical studies suggest. (c) Tractography results for 6 subjects viewed from an axial perspective in the template space. Here we show the consistent nature of the custom protocol in modeling of this complex pathway. In each of the subjects the U-shaped fanning bundle is clearly seen with individual difference in the amount of branching in the prefrontal cortex.

The creation of this region allowed the possibility of filtering out false-projections along the inferior longitudinal fasciculi and projections going medially toward the caudate, striatal areas, and anterior cingulate cortex. Additionally, we included bilateral uncinate fasciculus regions from the Johns Hopkins white matter atlas available through DSI-Studio as a “seed region” and Brodmann’s areas 11 and 47 as an “end” region. The tractography settings included the following settings: seed count = 500,000, threshold index = QA, FA threshold = 0.0, seed plan = “0”, initial direction=“0”, interpolation= “0”, step size = “0”, turning angle = “55”, smoothing = 0.6, min length = 0, max length = 150.

The tractography protocol was repeated for the left and right hemisphere separately by using the left or right uncinate “seed” region in each iteration. In order to quantify the integrity of the resulting streamlines we used the model-free measure of normalized quantitative anisotropy (nQA) in order to quantify the degree of diffusion along principle orientations on an ODF. Although this measure isn’t directly comparable to FA, it is possible that it more accurately represents the degree of diffusion among principle axonal directions within a single voxel rather than a singular tensor-based measure that does not account for multiple orientations. Its calculation is described in Yeh et al., 2013 and it captures the degree of anisotropy among major fiber populations within voxel accounting for the isotropic component and is normalized on a scale from 0-1. Together, these methods allowed for consistent modeling of the UF in a standardized way and unbiased by experimenter manual intervention.

### Statistical Analysis

All statistical analyses were conducted using Prism GraphPad 7 and R-studio. We conducted all correlational analysis in Prism 7 computing two-tailed Pearson correlation coefficients with 95% confidence intervals. Multiple linear regressions were used to access the influence of Beck Depression Inventory scores on relationships between behavior, fMRI, and diffusion measures and were conducted in R-studio. Mediation analysis was done using the *mediate* package in R with bootstrapped confidence intervals and 10,000 simulations. Statistical values were considered significant at alpha level of 0.05, which appropriately controlled for Type I error.

## Acknowledgements

We would like to acknowledge Ms. Jessica Noche, Ms. Amanda Chun, and Ms. Elizabeth Murray for their help with data collection and Dr. Zachariah Reagh and Mr. Jared Roberts (posthumously) for their assistance with analysis of the functional MRI data.

